# Virulence genes are a signature of the microbiome in the colorectal tumor microenvironment

**DOI:** 10.1101/009431

**Authors:** Michael B Burns, Joshua Lynch, Timothy K Starr, Dan Knights, Ran Blekhman

## Abstract

**Background:** The human gut microbiome is associated with the development of colon cancer, and recent studies have found changes in the composition of the microbial communities in cancer patients compared to healthy controls. However, host-bacteria interactions are mainly expected to occur in the cancer microenvironment, whereas current studies primarily use stool samples to survey the microbiome. Here, we highlight the major shifts in the colorectal tumor microbiome relative to that of matched normal colon tissue from the same individual, allowing us to survey the microbial communities at the tumor microenvironment, and provides intrinsic control for environmental and host genetic effects on the microbiome.

**Results:** We characterized the microbiome in 44 primary tumor and 44 patient-matched normal colon tissues. We find that tumors harbor distinct microbial communities compared to nearby healthy tissue. Our results show increased microbial diversity at the tumor microenvironment, with changes in the abundances of commensal and pathogenic bacterial taxa, including *Fusobacterium* and *Providencia*. While *Fusobacteria* has previously been implicated in CRC, *Providencia* is a novel tumor-associated agent, and has several features that make it a potential cancer driver, including a strong immunogenic LPS and an ability to damage colorectal tissue. Additionally, we identified a significant enrichment of virulence-associated genes in the colorectal cancer microenvironment.

**Conclusions:** This work identifies bacterial taxa significantly correlated with colorectal cancer, including a novel finding of an elevated abundance of *Providencia* in the tumor microenvironment. We also describe several metabolic pathways and enzymes differentially present in the tumor associated microbiome, and show that the bacterial genes in the tumor microenvironment are enriched for virulence associated genes from the aggregate microbial community. This virulence enrichment indicates that the microbiome likely plays an active role in colorectal cancer development and/or progression. These reuslts provide a starting point for future prognostic and therapeutic research with the potential to improve patient outcomes.

## Background

Colorectal cancer (CRC) is the second most commonly diagnosed cancer in females and the third in males worldwide[1]. The microbial communities present in the intestinal tract have known associations with colon health, though until recently researchers were limited to the study of microbes that were amenable to *in vitro* culturing. As a result of recent advances in culture-independent measurements of microbial communities, we know that the human gut is host to roughly a thousand different bacterial species[2]. Other alterations of this bacterial community are correlated with host health, including diseases from diabetes and obesity to Crohn’s disease and arteriosclerosis[3]. The composition of the gut microbiome has a known association with colorectal cancer, although the direction of causality remains unknown[4–10]. A recent report demonstrated that analysis of the microbiome can be used as a pre-screening test for CRC that outperformed several current standard methods[11]. These analyses have identified significant shifts in the relative abundances of specific bacterial taxa in CRC cancer patients’ colon mucosa and stool microbiomes. For instance, bacteria in the genus *Fusobacteria* are enriched in some CRC patients’ microbiomes[7, 8, 10, 12]. *Fusobacteria* are thought to elicit a pro-inflammatory microenvironment around the tumor, driving tumor formation and/or progression[7]. More specifically, a recent study has demonstrated that the FadA protein, a virulence factor expressed by *Fusobacterium nucleatum*, can signal epithelial cells via E-cadherin, a cell-surface molecule important for CRC metastasis as well as a component of the WNT/β-catenin signaling pathway that is the most commonly mutated pathway in CRC[13]. Other cancer-associated bacterial taxa have been identified as well, including *E. coli* strain NC101 and *Bacteroides fragilis,* each with a proposed mechanism of interaction with colon cancer[14]. The species *Akkermansia mucinphila*, a bacterium with known associations in obesity, has also been implicated as a cancer-associated agent, with its mucin-degrading activity as a proposed mechanism to drive inflammation contributing to cancer genesis and/or progression[14].

In addition to defining the set of significant bacteria associated with CRC, several groups have used measurements of microbiome diversity to compare cancer patients to normal subects. There are distinct differences in these results that depend on the sources of the samples used to assess the microbiome (*e.g.* stool samples versus mucosa or tissue samples, longitudinal versus cross-sectional sampling). For instance, in a study that used stool samples to compare CRC patients to normal controls, the researchers showed a decreased alpha-diversity among the microbial communities found in the CRC patients’ stools compared to the control[15]. However, in a different study that used tissue samples from patients with colon adenomas and compared them to patient matched normal tissues, the alpha-diversity present at the site of the lesion was actually increased[16]. This finding was repeated in a study by Mira-Pascual, *et al.* who performed side-by-side analyses of tissue and stool samples. They found that stool samples in general had roughly twice the microbial diversity when compared to tissue-associated microbiomes, though when comparing only tissues, the tumor microbiome was still more diverse than the normal microbiome[17]. It is likely that this is a function of the stool samples harboring microbes from the entire colonic environment, including species that are not directly related to the tumor microenvironment, adding noise to the taxonomic results acquired from assessment of stool samples relative to direct measurements of tissues. These findings suggest that in order to detect differences specific to the cancer-associated microbiome, samples taken directly from the tumor microenvironment are preferable, at least at the initial characterization phase, to bulk stool samples, which are not likely to have the discriminatory power required to measure small, yet significant, effects[17].

The use of traditional case-control studies of the colon cancer microbiome makes it difficult to control for all of the external effects on the microbiome. For example, the composition of gut microbial communities is strongly affected by diet[18]. Host genetic variation is also expected to control variation in the gut microbiome[19], through differences in host immune response and other genetic mechanisms[20]. The large effects these factors have on microbiome composition are likely to confound traditional case-control studies. By using tumor and normal tissue samples taken from the same individual, our study controls for these variables internally, providing a more accurate view of the tumor-associated shifts in the microbiome.

In addition to measuring bacterial taxa levels in colon cancer, it is also important to take into account the associated factors such as host genetics and gene expression as well as the microenvironmental metabolome. Independent research groups have attempted to uncover pertinent alterations in these factors and how they correlate with cancer state[21–23]. Of note, analysis of the CRC-associated metabolome highlighted differences in the biochemical composition of cancer patients’ stools. CRC patients were found to have higher levels of some amino acids and alterations in the levels of some short chain fatty acids (SFCA) in their stools when compared to controls[21]. Butyrate, an SFCA with known anti-cancer properties, was depleted in CRC patient stool samples as were several genera of butyrate producing bacteria[24]. Our work continues this effort by expanding the analysis of the CRC-associated microbiome to include virtual metagenomic profiling of the enzymes and pathways present in the colorectal cancer microbiome, with specific attention paid to assessing the presence of known virulence-associated genes[25].

## Results

### Tumors harbor microbiomes disctict from those at normal tissues

We obtained patient-matched normal and tumor colon tissue samples from the University of Minnesota Biological Materials Procurement Network (BioNet) from 44 patients (all work was approved by the University of Minnesota Institutional Reivew Board, IRB protocol 1310E44403; see Additional file 1 for sample information). We assessed the microbiome associated with each sample by Illumina sequencing across the V5-V6 hypervariable regions of the 16S rRNA gene (see Methods section for details). This analysis showed variation in the bacterial phyla abundance when comparing the matched normal and tumor tissues (Figure 1A). This variability is consistent with previous reports and demonstrates that indeed, there is a cancer-associated signature in the tumor microbiome[6, 10, 15, 16, 26–28]. At the level of the phyla, each sample was dominated by firmicutes, bacteroidetes, and proteobacteria. There were clear and significant changes in these phyla between the normal and cancer states, with the tumors showing an enrichment of proteobacteria and a depletion of firmicutes and bacteroidetes (Figure 1B). Also consistent with previous reports, we saw an increase in the phylum fusobacteria in the tumor-associated microbiome (two-sided Wilxocon signed rank test q ≤ 0.1 after FDR correction for multiple tests)[7, 10, 12, 28].

**Figure 1.**
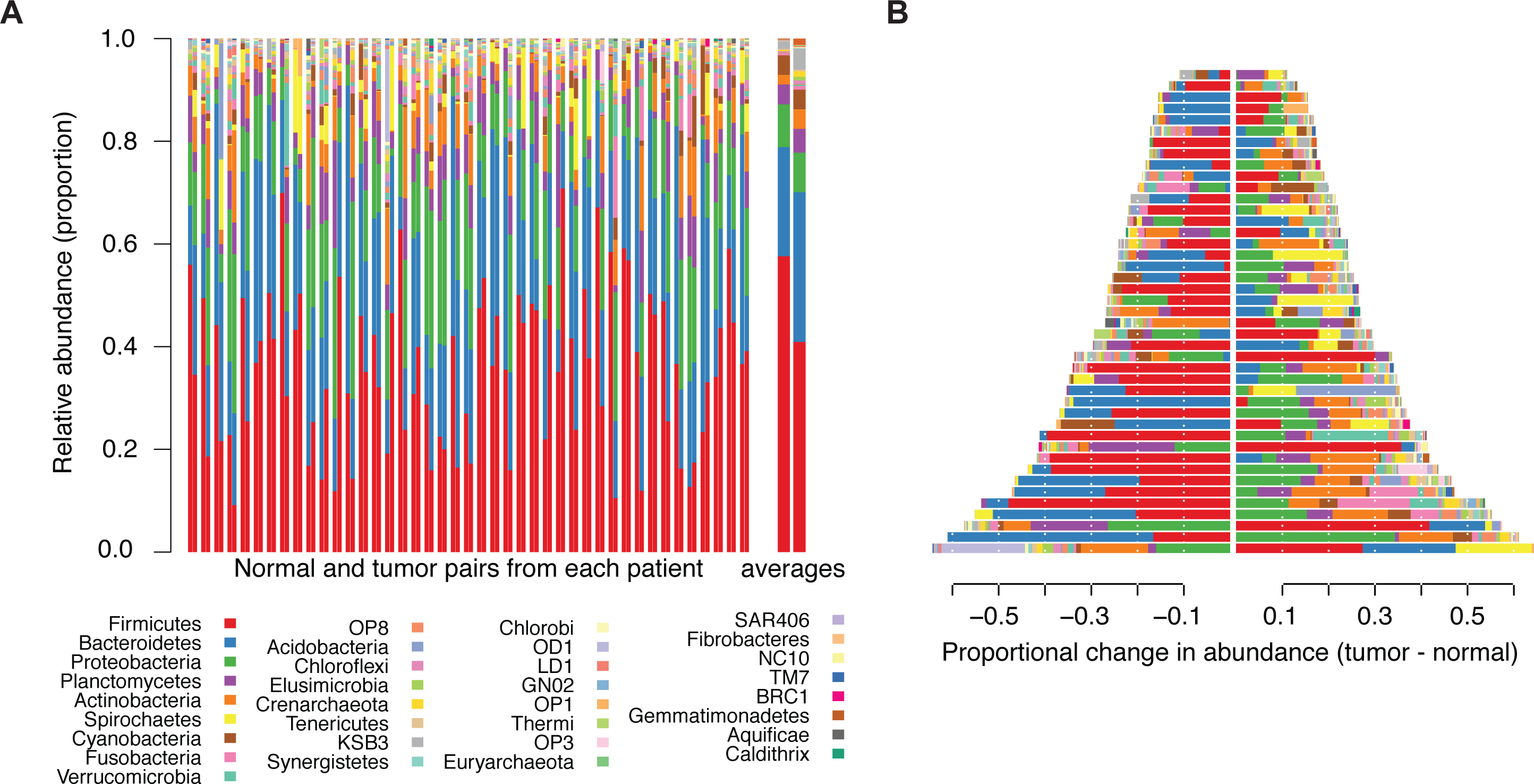
Differences in bacterial and archaeal phyla within the normal and colorectal cancer microbiomes. **(A)** Stacked bar plots indicating the proportional abundances of microbial phyla that are present at ≥1% in at least one sample. Columns are arranged as patient matched normal (left) and tumor (right) pairs, with the final, larger pair at the right representing the averages across all normals and tumors, repsectively. **(B)** The data presented in panel A, presented as the difference in abundance (tumor – normal) for each phylum, where a value of 0 would indicate no difference. Note that the legend is common to panels A and B.

When we assessed the differences at the level of operational taxonomic units (OTUs) we discovered numerous changes between the normal and tumor microbiomes with significant differences in the abundances of 19 different taxa (Wilxocon rank sum test q ≤ 0.1 after FDR correction). Of note, the tumors showed decreases in the abundances of several members of the order chlostridales, namely, lachnospiraceae, ruminococcaceae, and *Faecalibacterium prausnitzii*, as well as several members of the order bacteroidales, including *Bacteroides*, rikenellaceae, and *Bacteroides uniformis* (Figure 2 and Figure 3A). Taxa that were enriched in the tumor microbiomes included *Fusobacteria* and several proteobacteria genera including *Candidatus Portiera* and *Providencia* (Figure 2 and Figure 3A). Both *Fusobacterium* and *Providencia* are known pathogens, and when a correlation network is generated, it is clear that there are correlated abundance changes in the microbiome as a function of their presence (Figure 3B).

**Figure 2.**
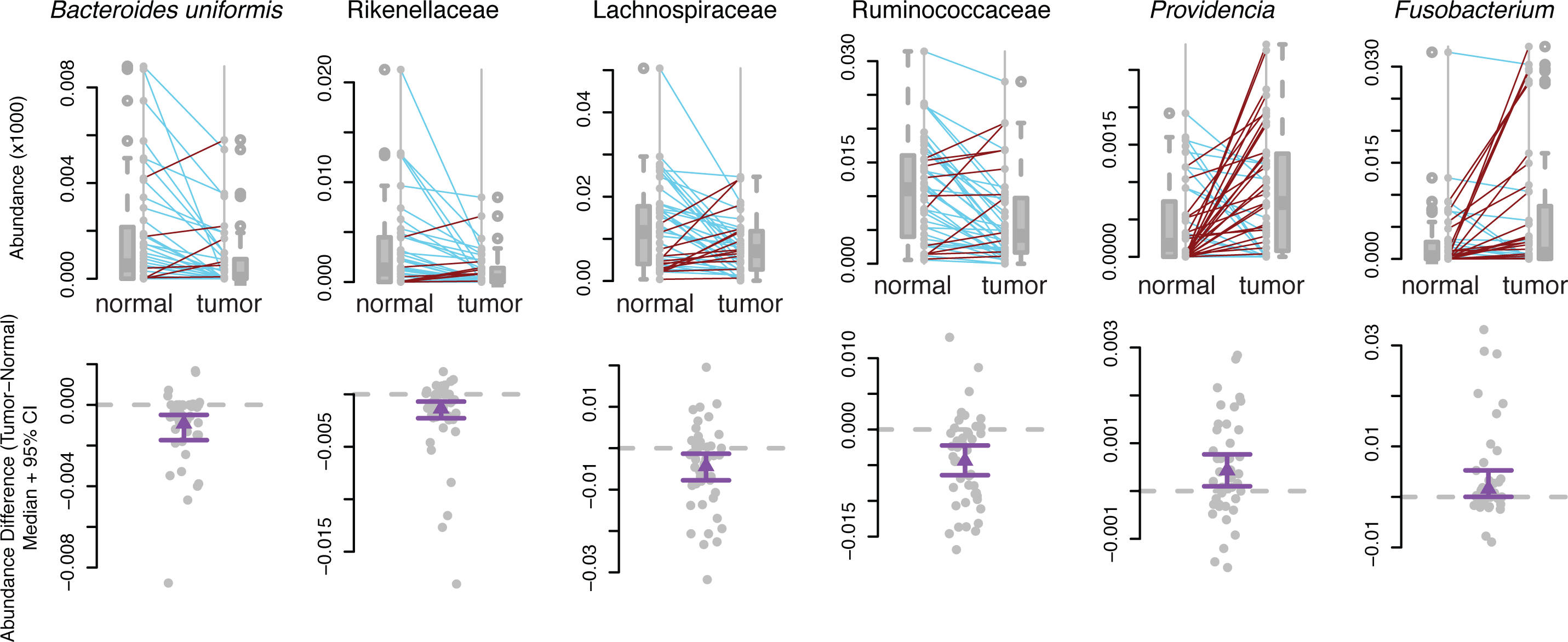
Differentially abundant OTUs between matched normal and colorectal cancer microbiomes. Boxplots with corresponding paired dotplots indicating the relative abundances of several differentially abundant OTUs. Lines connect the normal sample OTU abundance value (on left) to the tumor microbiome sample (on right). Line colors indicate the directionality of the abundance change (blue indicates a decreased aundance in the tumor relative to the normal, while red indicates an increased aundance in the tumor relative to the normal). These data are also each presented below each paired plot with the difference between tumor and normal abundance shown as grey dots. The purple line displayes the mean with a 95% confidence interval. A dashed horizontal grey line is plotted at 0.

**Figure 3.**
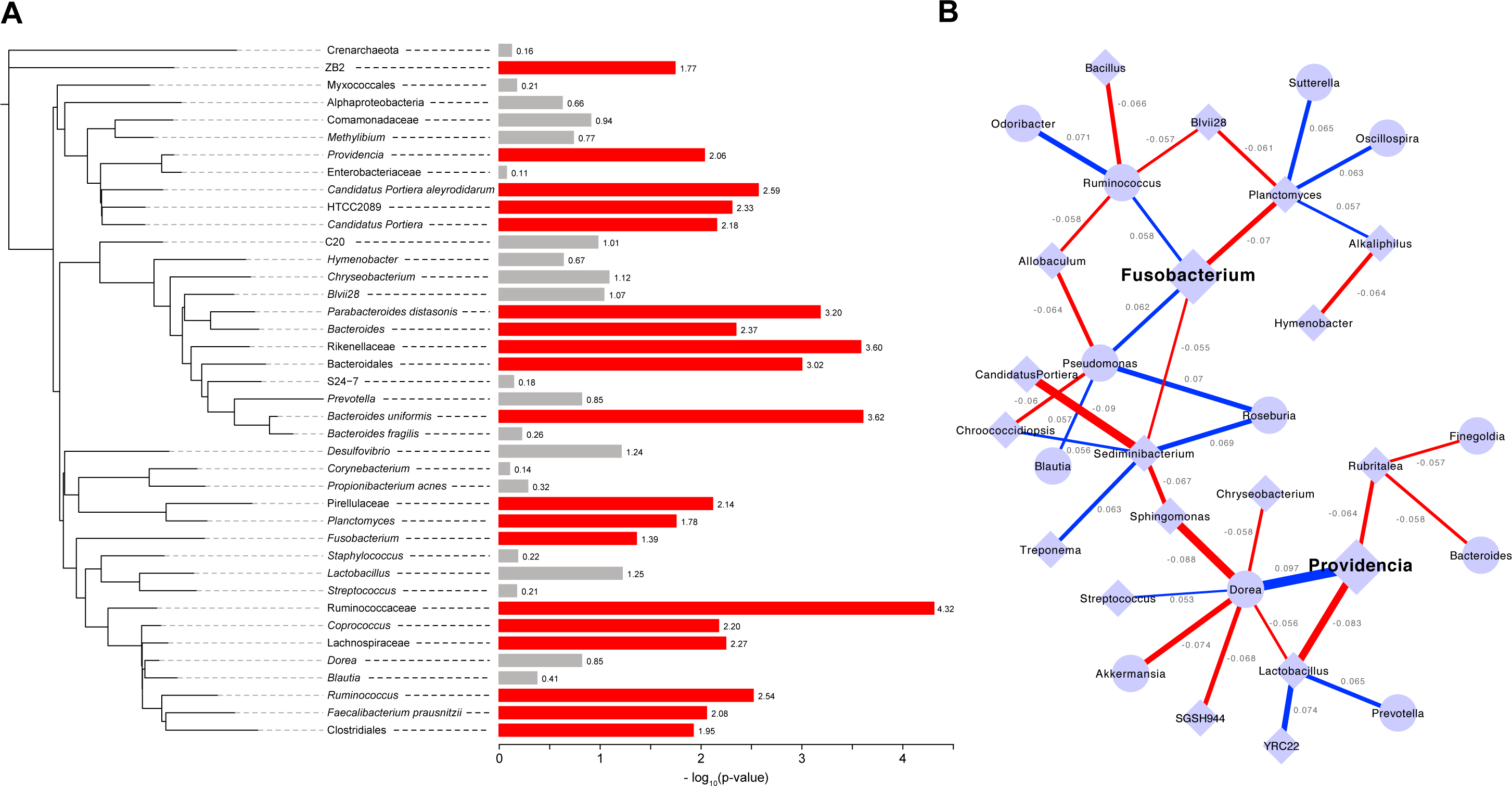
Relationships among the taxa found in colorectal cancer patients’ microbiomes. **(A)** Phylogenetic tree depicting the relatedness of the bacterial taxa present (> 0.1% of total) within 50% or more of the samples. The bars to the right indicate the –log_10_(p-value) from the Wilcoxon rank sum test to determine differential abundace between the normal and tumor microbiome. Red bars indicate significance at 10% FDR, while gray bars indicate that the specific taxon did not reach significance. **(B)** Correlation network showing the relationship among the abundaces of genera with absolute values of 0.05 or more and statistical significance (pseudo p-value <0.05). Edges indicate correlations: the edge thickness represents the magnitude and the color represents the sign (blue is positive correlation, red is negative correlation). Each node is a microbial genus where diamond shaped nodes indicate a higher average abundance and circular nodes indicate a lower average abundance in the tumor microbiome compared to normal.

### CRC-associated microbiome diversity

We calculated alpha-diversity using a variety of metrics within each of the samples using QIIME[29]. Alpha diversity metrics that account for phylogenetic relationships between the OTUs show that the tumor microbiomes exhibited higher alpha diversity than those of the normal, patient-matched microbiomes (p = 0.029 by two-sided Wilcoxon signed rank test). This is also true when using alternative measures of diversity such as the Shannon’s index or the Inverse Simpson’s (p = 0.020 and 0.024, respectively, by Wilcoxon rank sum test) (Figure 4).

**Figure 4.**
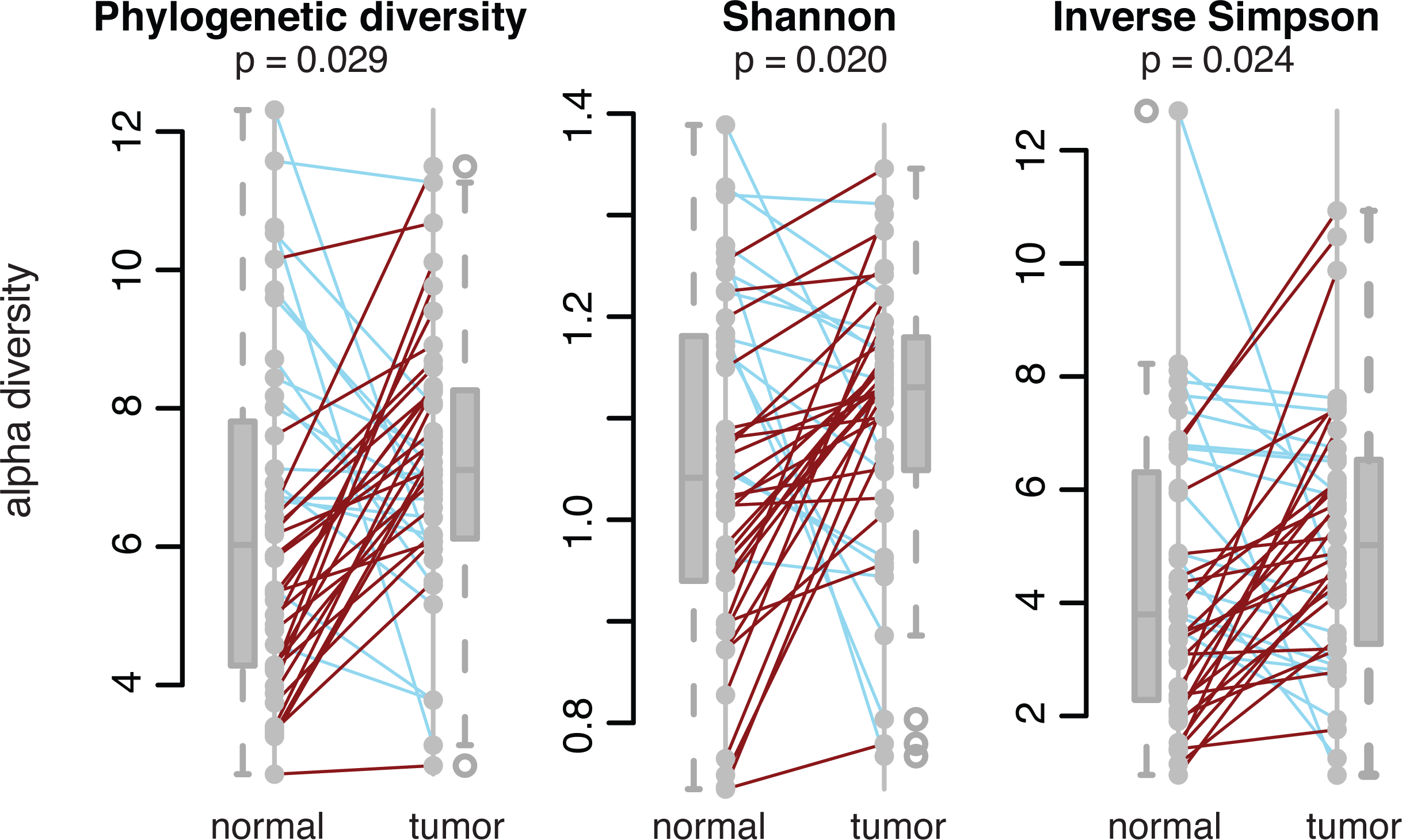
Microbial diversity within normal and tumor-associated microbiomes. Paired line plots shows the phylogenetic diversity, Shannon’s Index, and Inverse Simpson’s Index (alpha diversity metrics) for the microbiomes associated with normal and patient-matched tumor samples. The colors of the lines represent the direction of the change for each matched pair (blue lines indicate a decrease in diversity from normal to tumor, while red lines indicate an increase in diversity from the normal to tumor). P-values were calculated using a two-sided Wilcoxon signed rank test.

### Variation in the functional pathways and enzymes in the tumor microbiome

Using the PICRUSt (Phylogenetic Investigation of Communities by Reconstruction of Unobserved States) pipleine, we contructed a virtual metagenome for each of the samples’ microbiomes[25]. The KEGG database was used as a reference to determine the abundances of metabolic pathways and enzymes within the virtual metagenomes [30, 31]. As with the bacterial phyla, we saw significant variation in the functional pathways represented within each of the sampled microbiomes (Figure 5A), though, as expected from previous studies, we find that the variability in phylum abundances is far greater that the variability in the functional pathways (Figure 5B)[2, 32].

**Figure 5.**
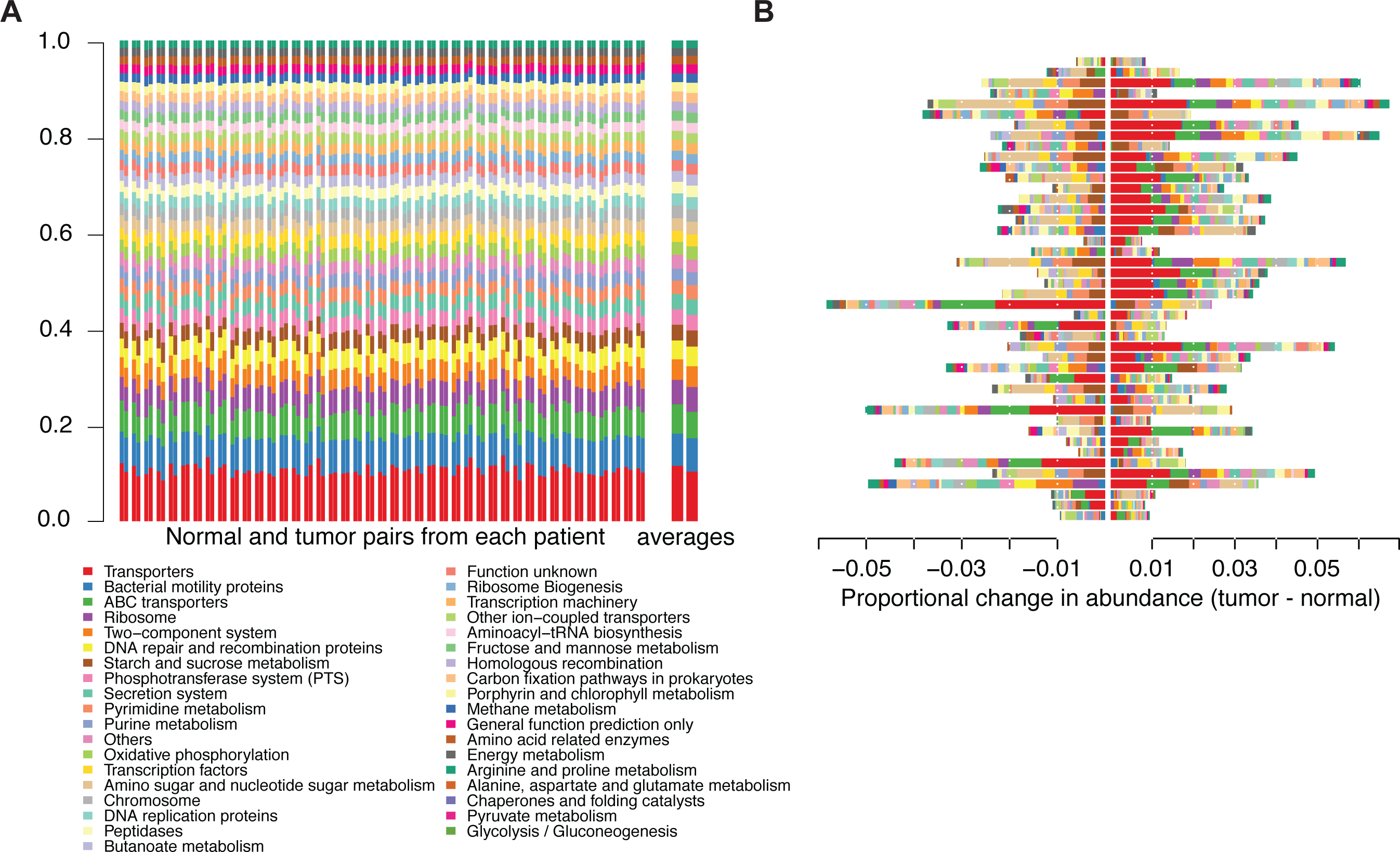
Differences in metabolic (KEGG) pathways within the normal and colorectal cancer microbiomes. **(A)** Stacked bar plots indicating the proportional pathway abundances within each sample that are present at ≥1% in at least one sample. Columns are arranged as patient matched normal (left) and tumor (right) pairs, with the final, larger pair at the right representing the averages across all normals and tumors, repsectively. **(B)** The data presented in panel A, showing the difference in abundance (tumor – normal) for each pathway, where a value of 0 would indicate no difference. Note that the legend is common to panels A and B.

These observations suggest that there is substatial functional redundancy across the phyla, in that while there might be differences in the taxa represented, on a functional level, many of them perform the same role in the gut. The patient-by-patient variability in phyla does not perfectly correspond to that seen at the functional pathway level, as expected, though analysis at the level of enzymes and pathways provides insights that analyses of the taxa alone may miss, due to the decreased between-sample variation. In general, the differences seen at the pathway level are roughly an order of magnitude less than the differences seen at the level of the phylum level (compare Figure 1B and Figure 5B). Although the pathway differences are smaller than those at the level of the phylum, there remain statistically significant, physiologically relevant changes between the normal and tumor metagenomes.

Numerous pathways (20, as defined by KEGG, level 3) were found to be differentially abundant between the tumor and normal tissue. Alanine, aspartate, and glutamate metabolism, DNA replication proteins, and starch and sucrose metabolism were significantly depleted in the tumor microbiome (q ≤ 0.01 for each by two-sided Wilcoxon signed rank test after FDR correction; Figure 6A). Conversely, secretion system, two-component system, and bacterial motility protein pathways were significantly enriched in the tumor microbiome (q ≤ 0.04 for each pathway by two-sided Wilcoxon signed rank test after FDR correction; Figure 6A). To more closely examine the variation in the microbiome as a function of cancer status, we also assessed the virtual metagenome at the level of specific enzyme abundances. Each of these enzymes were annotated with information regarding known virulence associations from MVirDB[33]. There was a clear enrichement in the tumor microbiome of enzymes related to microbial virulence when including all possible virulence categories (p = 0.0046 by Fisher’s exact test) (Figure 6B, C). When assessing enrichment for virulence related genes by functional category in MVirDB, the tumors were significantly enriched for genes encoding general virulence proteins (p = 5.8 × 10^−5^, by Fisher’s exact test). Genes encoding bacterial toxins were found at higher abundance in the tumor, but the enrichment was not statistically significant (p = 0.17 by Fisher’s exact test), likely due to low total gene counts for some categories (*e.g.* protein toxins).

**Figure 6.**
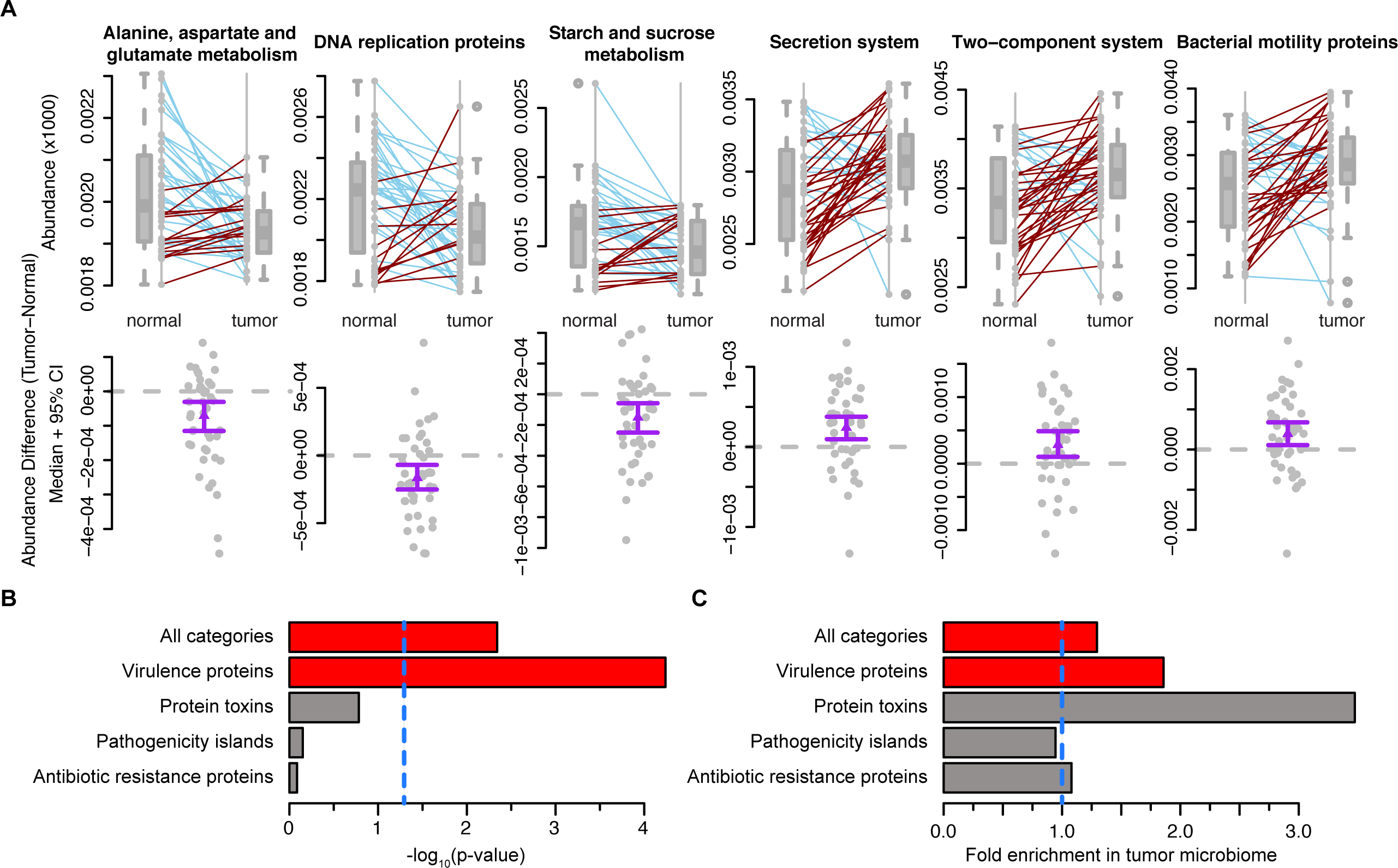
Differentially abundant pathways and enzyme classes between matched normal and colorectal tissue microbiomes. **(A)** Boxplots with corresponding paired dotplots indicating the relative abundances of several differentially abundant pathways. Lines connect the normal microbiome pathway abundance value (left) to the tumor microbiome value (right). Line colors indicate the directionality of the abundance change (blue for a decreased aundance in the tumor relative to the normal and red for an increased aundance in the tumor relative to the normal. These data are also each presented below each paired plot with the tumor abundance – normal abundance shown as grey dots. The purple line with an arrow represents the 95% CI in the indicated direction from the mean. Values at 0 (grey dotted lines) represent no change between normal and tumor. **(B)** Barchart showing the p-values (-log_10_ transformed) obtained from Fisher’s exact test used to determine virulence category enrichment in the tumor-associated microbiome on the x-axis with the gene categories labeled on the y-axis. Red bars indicate significance by Fisher’s exact test (p <0.005) and gray bars indicate no statistical significance. The blue, dashed line indicates the standard signifiance cut off of p = 0.05. **(C)** Barchart from the analysis in panel B, demonstrating the fold-enrichment of virulence protein-encoding genes in the tumor-associate microbiome. The x-axis is the fold enrichment of the different virulence enzyme classes within the tumor microbiome relative to the normal microbiome. The vertical, blue, dotted line placed at 1 indicates the point where there is no difference between the normal and tumor microbiomes.

## Discussion

At the phylum level, the differences seen between the normal and tumor tissue associated microbiomes are consistent with many previous reports[6, 10, 15, 16, 26– 28]. When assessing the data using information that accounts for more fine-grained detail with respect to taxonomy, we have made several important findings. Two of the genera we found to be enriched in the tumor microbiome, *Providencia* and *Fusobacteria*, are known to be pathogenic; *Fusobacteria* has been implicated previously in CRC[7, 10, 12, 28].

Species belonging to the genus *Providencia* have been implicated as infectious agents causing urinary tract infections, ocular infections, and gastroenteritis[34–36]. In addition, it is a genus in which some sub-strains having acquired resistance to commonly used antibiotics[36–38]. *Fusobacterium* is a genus that encompass species known to be pathogenic in humans; they are obligate anaerobes, with known sites of infection typically in the gastrointestinal tract[39]. The finding that these particular genera are prevalent in the tumor microenvironment implies several alternative hypotheses. One possibility is that these bacteria are causative in oncogenesis or tumor progression; another possibility is that these species are being enriched as the tumor has formed a niche that favors these bacteria. In the case of *Fusobacteria*, the results from several different studies, both correlative and mechanistic, indicate that it is likely a cancer driver[7, 28, 40]. In the case of *Providencia*, there are as yet no definitive studies that implicate this genus as a contributor to colorectal cancer. The discovery of *Providencia* in the tumor microbiome is interesting as, similar to *Fusobacteria*, it encodes a potent, immunogenic lipopolysaccharide (LPS) and can disrupt the epithelial membrane in the intestines, though the mechanism by which this is accomplished is still unclear[35, 41–43]. These factors manifest phenotypically as gastroenteritis, though with our discovery of its association with the cancer microenvironment, it is a promising candidate cancer-promoting pathogen.

From a diagnostic and therapeutic perspective, assessing the CRC-associated microbiome as a function of the different taxa is an eminently worthwhile endeavour as it is the logical location to look for specific taxa that could be biomarkers and/or targets for intervention in CRC. However, it is possible that the search for specific taxa might miss the larger perspective. For instance, as described above, *Fusobacteria* and *Providencia* share some important phenotypic characteristics – potent, immunogenic LPS and the ability to damage colorectal tissue. These similarities might be better assessed using metagenomic or metatranscriptomic approaches, virtual or otherwise, as these key features are undoubtedly reflected in the genes that these particular bacteria encode. This report is the first to highlight the feasability of such an approach by showing the striking enrichment of virulence genes in the tumor-associated microbiome. The fact that virulence proteins are enriched in the tumor-associated microbiome lends support to the hypothesis that the microbiome is an active contributor to colorectal cancer, rather than the result of a bystander phenomenon. It is important to note that this clear enrichment is likely underestimated because MVirDB, while expansive, does not currently encompass all known virulence genes in the microbiome, and, as the field of medical microbial genomics advances, new virulence genes will undoubtedly be discovered. For instance, the FadA protein from *Fusobacterium nucleatum* has been reported as a critical virulence factor, yet as it is a recent report, this finding has not yet made its way into MVirDB as of this submission[33][40].

## Conclusions

It is clear that there are numerous taxa in the CRC microbiome that are correlated with the disease. Here, in addition to the previously reported genus *Fusobacterium*, we report the discovery of another genus with similar pathogenic features, *Providencia*. This manuscript also presents an analysis that incorporates information at the functional (*e.g.* virulence potential) level to assess differences between the normal and cancer-associated microbiomes. It is important to note that these two approaches (taxonomy-based and function-based) are complementary and can provide insights into the pathogenic potential of the taxa and genes found in the microbiome of the tumor microenvironment. Our work demonstrates that utilizing both approaches can provide researchers with specific taxa as biomarkers and/or therapeutic targets while also looking globally at the pathogenic potential of the microbiome and showing a clear enrichment of virulence-associated microbial genes present in the CRC-microbiome. As with the bacterial genera associated with the disease, these virulence genes may provide researchers and clinicians with tergets for therapeutic intervention to improve patient outcomes.

## Methods

### Tissue samples and DNA isolation

We used 88 tissue samples from 44 individuals, with one tumor and one normal sample from each individual (see Additional file 1 for detailed information on the samples used in this study). Total DNA and RNA were isolated from flash-frozen colon tissue samples and their associated microbiomes by adapting an established protocol[44]. Briefly, approximately 100 mg of flash-frozen tissue were physically disrupted by placing the tissue in 1mL of Qiazol and sonicting in a heated (65°C) ultasonic water bath for 1-2 hours. DNA was purified from the lysate using the Qiagen All-prep kit.

### 16S rRNA sequencing

DNA isolated from colon samples was quantified by qPCR, and the V5-V6 regions of the 16S rRNA gene were PCR amplified with the addition of barcodes for multiplexing. The forward and reverse primers were the V5F and V6R sets from Cai, *et al.* [45]. The barcoded amplicons were pooled and Illumina adapters were ligated to the reads. A single lane on an Illumina MiSeq instrument was used (250 cycles, paired-end) to generate 16S rRNA gene sequences.

### Sequence Analysis

The sequence data contained approximately 10.7 million total reads passing quailty filtering total, with a mean value of 121,470 quality reads per sample. The forward and reverse read pairs were merged using the USEARCH v7 program ‘fastq_mergepairs’, allowing stagger, with no mismatches allowed[46]. Merged reads were quality trimmed, again using USEARCH, to truncate reads at any quality scores of 20 or less. The fasta sequence headers were renamed using a custom script to conform to QIIME standards.

The merged and filtered reads were used to pick OTUs using with QIIME v1.7.0 using ‘pick_otus.py’, with the closed-reference usesearch_ref OTU picking protocol against the Greengenes database (August 2013 release) at 97% similarity[29, 47, 48]. Reverse read matching was enabled, while reference based chimera calling was disabled. Rarefaction was performed on the otu table at 5000 reads prior to subsequent analyses.

The final OTU table was used to generate a phylogenetic tree by first filtering the taxa to include only those that were present at at least 0.1% in at least 50 of samples. This set was used as a starting point to trim the Greengenes database (August 2013 release) provided tree file (97_otus_unannotated.tree) using a custom pipeline (Sycamore – available at https://github.com/almlab/sycamore) from the Alm laboratory at MIT. The output of this pipeline was visualized with the Interactive Tree of Life[48, 49]. See Additional file 2 for the OTU table used in this study.

We used a linear model to correct for several patient and tumor covariates, individually as well as in combination, including patient age, sex, tumor stage, and tumor site. None of these factors, alone or in combination, were found to have a significant impact in this sample set. We note that microsatellite instable/microsatellite stable (MSI/MSS) statuses were only avialble for 13 of the 44 patients.

Correlation analsis was performed using SparCC, available at https://bitbucket.org/yonatanf/sparcc from Jonathan Friedman at MIT, on the complete OTU table collapsed to the genus level[50]. Pseudo p-values were inferred using 100 randomized sets. Correlations with pseudo p-values ≤0.05 that were within two degrees of separation from *Providencia* or *Fusobacterium* with absolute correlations of 0.05 or more were visualized using Cytoscape v3.1.0.

The PICRUSt v1.0.0 pipeline was used to generate a virtual metagenome using the OTU table generated in the previous analyses by QIIME[25, 29, 47]. Pathways and enzymes were assigned using the KEGG database options built into the pipeline. Virulence genes were identified by mapping the data in the PICRUSt enzyme abundance table to MVirDB using the uniprot database file, idmapping.dat, available from http://www.uniprot.org, as a key. See Additional files 3 and 4 for the metabolic enzyme and pathway abundance tables, respectively.

## Authors’ contributions

MBB and RB concieved of the study and study design. MBB carried out DNA extraction, sequence analyses, figure generation, and manuscript preparation. RB provided guidiance with statistical analyses and assisted in figure generation. JL assisted in statistical analyses. TKS provided support with the experimental design related to patient sample inclusion and data interpretation. DK provided guidance with respect to the PICRUSt analysis and experimental design. All authors contributed to manuscript revisions, read, and approved of the final version.

## Acknowledgements

The authors thank the members of the Blekhman lab for useful doscussions, the Huttenhower lab at Harvard University for providing a publicly available Galaxy server for use with PICRUSt software, and the Alm lab at MIT for making their Sycamore pipeline publicly available at GitHub. This work was carried out, in part, using computing resources at the Minnesota Supercomputing Institute.

## Additional files

**Additional file 1 – Sample and patient clinical information and metadata**

A tab-delimited file containing a sample map that containing patient and tumor clinical information as well as the specific sample numbers that will allow proper identification of tumor and normal matched pairs.

**Additional file 2 – Unfiltered OTU table**

A tab-delimited file containing the unfiltered OTU table with proportional abundances of the taxa output from the QIIME pipeline.

**Additional file 3 – Enzyme abundances**

A tab-delimited file containing the unfiltered proportional abundances of enzymes detected in the virtual metagenomes of each sample using PICRUSt.

**Additional file 4 – Pathway abundances**

A tab-delimited file containing the unfiltered level 3 KEGG pathway proportional abundances for each sample generated using PICRUSt.

## References

1. Jemal A, Bray F, Center MM, Ferlay J, Ward E, Forman D: Global cancer statistics. CA Cancer J Clin 2011, 61:69–90.

2. Consortium THMP: Structure, function and diversity of the healthy human microbiome. Nature 2012, 486:207–214.

3. Jones ML, Ganopolsky JG, Martoni CJ, Labbé A, Prakash S: Emerging science of the human microbiome. Gut Microbes 2014, 5.

4. Konstantinov SR, Kuipers EJ, Peppelenbosch MP: Functional genomic analyses of the gut microbiota for CRC screening. Nat Rev Gastroenterol Hepatol 2013, 10:741–745.

5. Sobhani I, Tap J, Roudot-Thoraval F, Roperch JP, Letulle S, Langella P, Corthier G, Tran Van Nhieu J, Furet JP: Microbial dysbiosis in colorectal cancer (CRC) patients. PloS One 2011, 6:e16393.

6. Bonnet M, Buc E, Sauvanet P, Darcha C, Dubois D, Pereira B, Déchelotte P, Bonnet R, Pezet D, Darfeuille-Michaud A: Colonization of the human gut by E. coli and colorectal cancer risk. Clin Cancer Res Off J Am Assoc Cancer Res 2014, 20:859–867.

7. Castellarin M, Warren RL, Freeman JD, Dreolini L, Krzywinski M, Strauss J, Barnes R, Watson P, Allen-Vercoe E, Moore RA, Holt RA: Fusobacterium nucleatum infection is prevalent in human colorectal carcinoma. Genome Res 2012, 22:299–306.

8. Tahara T, Yamamoto E, Suzuki H, Maruyama R, Chung W, Garriga J, Jelinek J, Yamano H, Sugai T, An B, Shureiqi I, Toyota M, Kondo Y, Estécio MRH, Issa J-PJ: Fusobacterium in colonic flora and molecular features of colorectal carcinoma. Cancer Res 2014, 74:1311–1318.

9. Chen W, Liu F, Ling Z, Tong X, Xiang C: Human intestinal lumen and mucosa-associated microbiota in patients with colorectal cancer. PloS One 2012, 7:e39743.

10. Kostic AD, Gevers D, Pedamallu CS, Michaud M, Duke F, Earl AM, Ojesina AI, Jung J, Bass AJ, Tabernero J, Baselga J, Liu C, Shivdasani RA, Ogino S, Birren BW, Huttenhower C, Garrett WS, Meyerson M: Genomic analysis identifies association of Fusobacterium with colorectal carcinoma. Genome Res 2012, 22:292–298.

11. Zackular JP, Rogers MAM, Ruffin MT, Schloss PD: The Human Gut Microbiome as a Screening Tool for Colorectal Cancer. Cancer Prev Res (Phila Pa) 2014.

12. Marchesi JR, Dutilh BE, Hall N, Peters WHM, Roelofs R, Boleij A, Tjalsma H: Towards the human colorectal cancer microbiome. PloS One 2011, 6:e20447.

13. Comprehensive Molecular Characterization of Human Colon and Rectal Cancer. Nature 2012, 487:330–337.

14. Irrazábal T, Belcheva A, Girardin SE, Martin A, Philpott DJ: The multifaceted role of the intestinal microbiota in colon cancer. Mol Cell 2014, 54:309–320.

15. Ahn J, Sinha R, Pei Z, Dominianni C, Wu J, Shi J, Goedert JJ, Hayes RB, Yang L: Human gut microbiome and risk for colorectal cancer. J Natl Cancer Inst 2013, 105:1907–1911.

16. Shen XJ, Rawls JF, Randall T, Burcal L, Mpande CN, Jenkins N, Jovov B, Abdo Z, Sandler RS, Keku TO: Molecular characterization of mucosal adherent bacteria and associations with colorectal adenomas. Gut Microbes 2010, 1:138– 147.

17. Mira-Pascual L, Cabrera-Rubio R, Ocon S, Costales P, Parra A, Suarez A, Moris F, Rodrigo L, Mira A, Collado MC: Microbial mucosal colonic shifts associated with the development of colorectal cancer reveal the presence of different bacterial and archaeal biomarkers. J Gastroenterol 2014.

18. David LA, Maurice CF, Carmody RN, Gootenberg DB, Button JE, Wolfe BE, Ling AV, Devlin AS, Varma Y, Fischbach MA, Biddinger SB, Dutton RJ, Turnbaugh PJ: Diet rapidly and reproducibly alters the human gut microbiome. Nature 2014, 505:559–563.

19. Cullender TC, Chassaing B, Janzon A, Kumar K, Muller CE, Werner JJ, Angenent LT, Bell ME, Hay AG, Peterson DA, Walter J, Vijay-Kumar M, Gewirtz AT, Ley RE: Innate and adaptive immunity interact to quench microbiome flagellar motility in the gut. Cell Host Microbe 2013, 14:571–581.

20. Knights D, Lassen KG, Xavier RJ: Advances in inflammatory bowel disease pathogenesis: linking host genetics and the microbiome. Gut 2013, 62:1505–1510.

21. Weir TL, Manter DK, Sheflin AM, Barnett BA, Heuberger AL, Ryan EP: Stool Microbiome and Metabolome Differences between Colorectal Cancer Patients and Healthy Adults. PLoS ONE 2013, 8.

22. Warren RL, Freeman DJ, Pleasance S, Watson P, Moore RA, Cochrane K, Allen-Vercoe E, Holt RA: Co-occurrence of anaerobic bacteria in colorectal carcinomas. Microbiome 2013, 1:16.

23. Ohigashi S, Sudo K, Kobayashi D, Takahashi O, Takahashi T, Asahara T, Nomoto K, Onodera H: Changes of the intestinal microbiota, short chain fatty acids, and fecal pH in patients with colorectal cancer. Dig Dis Sci 2013, 58:1717– 1726.

24. Weaver GA, Krause JA, Miller TL, Wolin MJ: Short chain fatty acid distributions of enema samples from a sigmoidoscopy population: an association of high acetate and low butyrate ratios with adenomatous polyps and colon cancer. Gut 1988, 29:1539–1543.

25. Langille MGI, Zaneveld J, Caporaso JG, McDonald D, Knights D, Reyes JA, Clemente JC, Burkepile DE, Vega Thurber RL, Knight R, Beiko RG, Huttenhower C: Predictive functional profiling of microbial communities using 16S rRNA marker gene sequences. Nat Biotechnol 2013, 31:814–821.

26. Arthur JC, Perez-Chanona E, Mühlbauer M, Tomkovich S, Uronis JM, Fan T-J, Campbell BJ, Abujamel T, Dogan B, Rogers AB, Rhodes JM, Stintzi A, Simpson KW, Hansen JJ, Keku TO, Fodor AA, Jobin C: Intestinal Inflammation Targets Cancer-Inducing Activity of the Microbiota. Science 2012, 338:120–123.

27. Wu N, Yang X, Zhang R, Li J, Xiao X, Hu Y, Chen Y, Yang F, Lu N, Wang Z, Luan C, Liu Y, Wang B, Xiang C, Wang Y, Zhao F, Gao GF, Wang S, Li L, Zhang H, Zhu B: Dysbiosis signature of fecal microbiota in colorectal cancer patients. Microb Ecol 2013, 66:462–470.

28. Kostic AD, Chun E, Robertson L, Glickman JN, Gallini CA, Michaud M, Clancy TE, Chung DC, Lochhead P, Hold GL, El-Omar EM, Brenner D, Fuchs CS, Meyerson M, Garrett WS: Fusobacterium nucleatum potentiates intestinal tumorigenesis and modulates the tumor-immune microenvironment. Cell Host Microbe 2013, 14:207–215.

29. Caporaso JG, Kuczynski J, Stombaugh J, Bittinger K, Bushman FD, Costello EK, Fierer N, Peña AG, Goodrich JK, Gordon JI, Huttley GA, Kelley ST, Knights D, Koenig JE, Ley RE, Lozupone CA, McDonald D, Muegge BD, Pirrung M, Reeder J, Sevinsky JR, Turnbaugh PJ, Walters WA, Widmann J, Yatsunenko T, Zaneveld J, Knight R: QIIME allows analysis of high-throughput community sequencing data. Nat Methods 2010, 7:335–336.

30. Kanehisa M, Goto S, Sato Y, Kawashima M, Furumichi M, Tanabe M: Data, information, knowledge and principle: back to metabolism in KEGG. Nucleic Acids Res 2014, 42(Database issue):D199–205.

31. Kanehisa M, Goto S: KEGG: kyoto encyclopedia of genes and genomes. Nucleic Acids Res 2000, 28:27–30.

32. Carroll IM, Chang Y-H, Park J, Sartor RB, Ringel Y: Luminal and mucosal-associated intestinal microbiota in patients with diarrhea-predominant irritable bowel syndrome. Gut Pathog 2010, 2:19.

33. Zhou CE, Smith J, Lam M, Zemla A, Dyer MD, Slezak T: MvirDB--a microbial database of protein toxins, virulence factors and antibiotic resistance genes for bio-defence applications. Nucleic Acids Res 2007, 35(Database issue):D391–394.

34. Koreishi AF, Schechter BA, Karp CL: Ocular infections caused by Providencia rettgeri. Ophthalmology 2006, 113:1463–1466.

35. Murata T, Iida T, Shiomi Y, Tagomori K, Akeda Y, Yanagihara I, Mushiake S, Ishiguro F, Honda T: A large outbreak of foodborne infection attributed to Providencia alcalifaciens. J Infect Dis 2001, 184:1050–1055.

36. Lau S-M, Peng M-Y, Chang F-Y: Resistance rates to commonly used antimicrobials among pathogens of both bacteremic and non-bacteremic community-acquired urinary tract infection. J Microbiol Immunol Infect Wei Mian Yu Gan Ran Za Zhi 2004, 37:185–191.

37. Lee H-W, Kang H-Y, Shin K-S, Kim J: Multidrug-resistant Providencia isolates carrying blaPER-1, blaVIM-2, and armA. J Microbiol Seoul Korea 2007, 45:272–274.

38. Luzzaro F, Mezzatesta M, Mugnaioli C, Perilli M, Stefani S, Amicosante G, Rossolini GM, Toniolo A: Trends in production of extended-spectrum beta-lactamases among enterobacteria of medical interest: report of the second Italian nationwide survey. J Clin Microbiol 2006, 44:1659–1664.

39. Allen-Vercoe E, Strauss J, Chadee K: Fusobacterium nucleatum: an emerging gut pathogen?. Gut Microbes 2011, 2:294–298.

40. Rubinstein MR, Wang X, Liu W, Hao Y, Cai G, Han YW: Fusobacterium nucleatum promotes colorectal carcinogenesis by modulating E-cadherin/β-catenin signaling via its FadA adhesin. Cell Host Microbe 2013, 14:195–206.

41. Asakura H, Momose Y, Ryu C-H, Kasuga F, Yamamoto S, Kumagai S, Igimi S: Providencia alcalifaciens causes barrier dysfunction and apoptosis in tissue cell culture: potent role of lipopolysaccharides on diarrheagenicity. Food Addit Amp Contam Part A 2013, 30:1459–1466.

42. Albert MJ, Alam K, Ansaruzzaman M, Islam MM, Rahman AS, Haider K, Bhuiyan NA, Nahar S, Ryan N, Montanaro J: Pathogenesis of Providencia alcalifaciens-induced diarrhea. Infect Immun 1992, 60:5017–5024.

43. Guth BE, Perrella E: Prevalence of invasive ability and other virulence-associated characteristics in Providencia alcalifaciens strains isolated in São Paulo, Brazil. J Med Microbiol 1996, 45:459–462.

44. Burns MB, Lackey L, Carpenter MA, Rathore A, Land AM, Leonard B, Refsland EW, Kotandeniya D, Tretyakova N, Nikas JB, Yee D, Temiz NA, Donohue DE, McDougle RM, Brown WL, Law EK, Harris RS: APOBEC3B is an enzymatic source of mutation in breast cancer. Nature 2013, 494:366–370.

45. Cai L, Ye L, Tong AHY, Lok S, Zhang T: Biased Diversity Metrics Revealed by Bacterial 16S Pyrotags Derived from Different Primer Sets. PLoS ONE 2013, 8.

46. Edgar RC: Search and clustering orders of magnitude faster than BLAST. Bioinforma Oxf Engl 2010, 26:2460–2461.

47. Navas-Molina JA, Peralta-Sánchez JM, González A, McMurdie PJ, Vázquez-Baeza Y, Xu Z, Ursell LK, Lauber C, Zhou H, Song SJ, Huntley J, Ackermann GL, Berg-Lyons D, Holmes S, Caporaso JG, Knight R: Advancing our understanding of the human microbiome using QIIME. Methods Enzymol 2013, 531:371–444.

48. DeSantis TZ, Hugenholtz P, Larsen N, Rojas M, Brodie EL, Keller K, Huber T, Dalevi D, Hu P, Andersen GL: Greengenes, a Chimera-Checked 16S rRNA Gene Database and Workbench Compatible with ARB. Appl Environ Microbiol 2006, 72:5069–5072.

49. Letunic I, Bork P: Interactive Tree Of Life v2: online annotation and display of phylogenetic trees made easy. Nucleic Acids Res 2011, 39(Web Server issue):W475–478.

50. Friedman J, Alm EJ: Inferring correlation networks from genomic survey data. PLoS Comput Biol 2012, 8:e1002687.

